# A deconvolution algorithm for multiecho functional MRI: Multiecho Sparse Paradigm Free Mapping

**DOI:** 10.1101/558288

**Authors:** César Caballero-Gaudes, Stefano Moia, Puja Panwar, Peter A. Bandettini, Javier Gonzalez-Castillo

**Affiliations:** Basque Center on Cognition, Brain and Language, San Sebastian, Spain.; Section on Functional Imaging Methods, Laboratory of Brain and Cognition, National Institute of Mental Health, National Institutes of Health, Bethesda, MD.; Functional MRI Core, National Institute of Mental Health, National Institutes of Health, Bethesda, MD.

**Keywords:** BOLD FMRI, MULTI-ECHO, DECONVOLUTION, SINGLE-TRIAL

## Abstract

This work introduces a novel algorithm for deconvolution of the BOLD signal in multiecho fMRI data: Multiecho Sparse Paradigm Free Mapping (ME-SPFM). Assuming a linear dependence of the BOLD percent signal change on the echo time (TE) and using sparsity-promoting regularized least squares estimation, ME-SPFM yields voxelwise time-varying estimates of the changes in the transverse relaxation 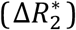 without prior knowledge of the timings of individual BOLD events. Our results in multi-echo fMRI data collected during a multi-task event-related paradigm at 3 Tesla demonstrate that the maps of 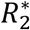 changes obtained with ME-SPFM at the times of the stimulus trials show high spatial and temporal concordance with the activation maps and BOLD signals obtained with standard model-based analysis. This method yields estimates of 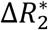 having physiologically plausible values. Owing to its ability to blindly detect events, ME-SPFM also enables us to map 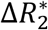 associated with spontaneous, transient BOLD responses occurring between trials. This framework is a step towards deciphering the dynamic nature of brain activity in naturalistic paradigms, resting-state or experimental paradigms with unknown timing of the BOLD events.

## INTRODUCTION

Task-based functional magnetic resonance imaging (fMRI) data is typically analyzed through the use of linear regression of BOLD signal change models on voxel time series. These regressors are defined assuming a linear model of the BOLD response as the convolution of a known activity with the hemodynamic response function (HRF). Recently, there has been an increasing interest in methods that enable to extract activation information without prior information of the timing of the BOLD events. Such methods can provide useful information about brain function in cases when insufficient knowledge about the neuronal activity driving the BOLD events is available, including naturalistic paradigms, resting state, and clinical conditions. In the absence of timing information, a potential approach is to estimate the activity-inducing signal underlying the BOLD responses; a process also known as deconvolution. Deconvolution allows detecting individual BOLD events (i.e. single trials) (Gaudes et al., 2011; Caballero-Gaudes et al., 2013), minimizing hemodynamic confounds in measures of functional connectivity (Gitelman et al., 2003; McLaren et al., 2012; Rangaprakash et al., 2018) and exploring time-varying activity of resting state fluctuations (Keilholz et al. 2017; Petridou et al., 2013; Karahanoğlu and Van de Ville, 2015, 2017). Here deconvolution is used to detect underlying events rather than to extract overlapping even-related hemodynamic responses when using a known timing (for example see Buckner et al., 1996; Goutte et al., 2000).

Deconvolution can also be understood as solving an inverse problem where the forward model is defined from the assumed hemodynamic model. If the deconvolution is performed with least squares estimation, estimates will be variable due to the high collinearity of the model. To overcome this, some type of regularization or prior information must be applied to the estimates of the activity-inducing signal. Initially, the deconvolution was done via empirical Bayesian estimators with Gaussian priors (Gitelman et al., 2003) or regularized least-squares estimators where the regularization term penalized the Euclidean norm (i.e. L_2_-norm) of the estimates (i.e. ridge regression) (Gaudes et al., 2011). Other approaches have employed sparsity-promoting regularized estimators based on the L_1_-norm or L_2,1_-norm of the estimates to improve the interpretability of the estimates, such as the Dantzig Selector, the Least Absolute Selection and Shrinkage Operator (LASSO) (Caballero-Gaudes et al., 2013, Khalidov et al., 2011) and a non-negative version of the fused LASSO (Hernandez-Garcia and Ulfarsson, 2011). The method of Total Activation incorporated spatio-temporal regularization terms based on generalized total variation and structured mixed L_2,1_-norms to improve the robustness of the deconvolution across neighboring voxels (Farouj et al., 2017; Karaganoglu et al., 2015). Structured mixed-norm regularization terms can also be used to account for variability in the shape of the assumed hemodynamic model (Gaudes et al., 2012). A nonparametric deconvolution method based on homomorphic filtering was proposed in Sreenivasan et al. (2015). Nonlinear regression methods using logistic functions have also been proposed to avoid assuming a linear model for the BOLD response (Bush and Cisler, 2013; Bush et al., 2015). Approaches using nonlinear state-space models (Riera et al., 2004), dynamic expectation maximization (Friston et al., 2018), generalized filtering (Friston et al., 2010) and its adaptation to a cubature Kalman filtering (Havlicek et al., 2011) have also been implemented to estimate the hidden activity-inducing signal and physiological parameters of the Balloon model of the BOLD response, which operate at a regional level to gain signal-to-noise ratio due to their higher complexity.

Relevant for the current work, all the aforementioned methods perform the deconvolution of fMRI data with one time series per voxel acquired at an echo time (TE). Acquisition at a single echo (1E) is commonly used for BOLD fMRI data, where the TE is usually chosen close to the average transverse relaxation parameter 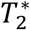 of the grey matter region of interest (Bandettini et al., 1994; Menon et al., 1993) to maximize the contrast-to-noise ratio of the signal. However, fMRI data can be alternatively acquired at multiple echo-times so that a weighted combination of the multiple echo signals can result in an enhancement in BOLD sensitivity, mainly in regions close to air-tissue boundaries that are prone to large signal dropouts and susceptibility distortions (Gowland and Bowtell, 2007; Poser et al., 2006; Posse et al., 1999; Posse, 2012). With multiecho fMRI (ME-fMRI) estimation of 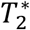 on a per-TR basis and voxel (i.e. a 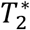-time series) is feasible, which can be used for subsequent analysis of task-related activity (Peltier and Noll, 2002) and functional connectivity (Wu et al., 2012, Power et al., 2018). Furthermore, ME-fMRI enables improved denoising of artefactual and confounding physiological signal fluctuations with dual-echo approaches (Bright and Murphy, 2013; Buur et al., 2009; Ing and Schwarzbauer, 2012) or multiecho independent component analysis (MEICA) (Kundu et al., 2012; 2013; 2017; Evans et al., 2015; Gonzalez-Castillo et al., 2016). Other denoising methods based on ME-fMRI acquisitions are discussed in Caballero-Gaudes and Reynolds (2017).

In this work, we propose a novel method for the temporal deconvolution of ME-fMRI data, named multiecho sparse paradigm free mapping (ME-SPFM). To our knowledge, no algorithm has been previously proposed for deconvolution of ME-fMRI data. Although previous approaches can be applied on ME-fMRI data after weighted combination of the multiple echo signals in a single dataset, the proposed approach directly operates with the multiple echo signals without combining them. Assuming a mono-exponential decay model of the gradient-echo signal, this method is able to estimate time-varying changes in the transverse relaxation rate 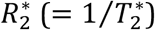, i.e. 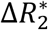, associated with single BOLD events without prior information of their timing. Using multiecho fMRI data acquired on 10 subjects (16 datasets) during an event-related paradigm including five distinct tasks (Gonzalez-Castillo et al., 2016), we demonstrate that the ME-SPFM algorithm considerably improves the accuracy of the deconvolution of individual BOLD events compared with its counterpart that operates in a single dataset or echo, namely sparse paradigm free mapping (hereafter denoted as 1E-SPFM) (Caballero-Gaudes et al., 2013). Furthermore, ME-SPFM yields voxel-wise quantitative estimates of 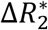 in interpretable units (s^−1^), which is relevant for functional analysis across different acquisition protocols and field strengths.

## METHODS

### Multiecho signal model

Assuming a mono-exponential decay model, the MR signal of a gradient echo acquisition in a voxel *x* at time *t* for an echo time TE_*k*_ can be approximated as

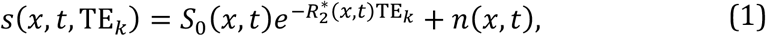

where *S*_0_(*x, t*) and 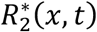 are the signal changes in the net magnetization *S*_0_ and the transverse relaxation rate 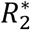 of the voxel *x* at time *t*, and *n*(*x, t*) is a noise term. Hereinafter, the noise term and the voxel index *x* are omitted for simplicity in the notation. Describing *S*_0_(*t*) and 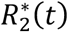 in terms of relative changes with respect to the average values in the voxel (Kundu et al., 2017), i.e. 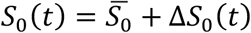 and 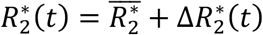, the MR signal can be written as

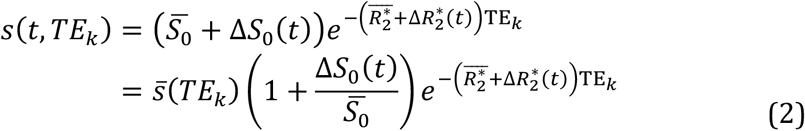

where the mean of the signal is 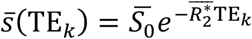. Typically, Δ*S*_0_(*t*) and 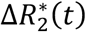 are considerably smaller than 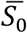 and 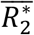, respectively. Hence, the last term in Eq. (2) can be approximated using a first-order Taylor approximation as 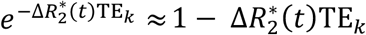. Substituting this term into Eq. (2) and defining 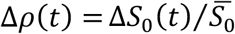, the MR signal can be approximated as

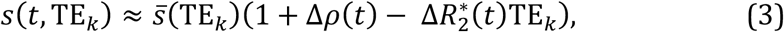

where the term resulting from the multiplication of small values of Δ*S*_0_(*t*) and 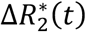 is neglected. Finally, signal percentage changes with respect to the mean of the signal, i.e. 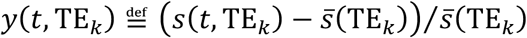, can be described as

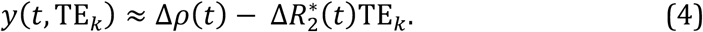

This signal model can be understood as a linear regression model in which the slope (i.e. dependent on the echo time TE_*k*_) captures the fluctuations related to 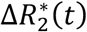, whereas the intercept captures the fluctuations related to Δ*S*_0_(*t*). In BOLD fMRI, signal changes related to 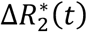 are more likely linked to neuronal processes than changes due to Δ*S*_0_(*t*), which are normally related to confounding effects such as motion or blood inflow.

Following the linear convolution model usually adopted in fMRI data analysis, let us also assume that changes in 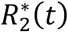 generating the BOLD response in the signal can be described as 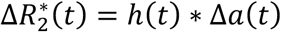, where Δ*a*(*t*) denotes an activity-inducing signal that is related to changes in neuronal activity, and *h*(*t*) is the hemodynamic response. Without lack of generality, we will assume that the shape of *h*(*t*) is independent of TE and also normalized to a peak amplitude equal to 1. Substituting in Eq. (4), signal percentage changes can then be approximated as

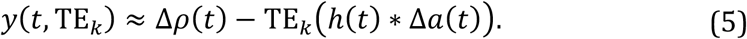

If signal changes related to variations in the net magnetization Δ*ρ*(*t*) are reduced during data preprocessing, the BOLD component of the signal can be approximated as

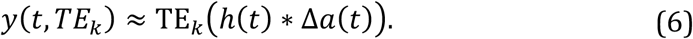

The continuous time MR signal is sampled every repetition time (TR), i.e. *t* = *n*TR, where *n* = 1,…,*N*, and *N* is the number of volumes acquired during the acquisition. In discrete time, the previous equations can be reformulated in matrix notation. We can define 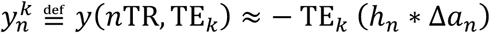, where 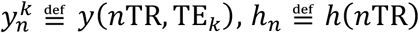, and 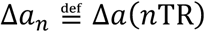. Gathering all time points as a vector, **y**_*k*_ = [*y*_1_,⋯,*y_N_*]^*T*^, we can write **y**_*k*_ ≈ −TE_*k*_**HΔ*a***, where **Δ*a*** ∈ ℝ^*N*^ is a column vector of length *N* that represents an activity inducing signal that is related to 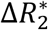, and **H** ∈ ℝ^***N×N***^ is a Toeplitz convolution matrix whose columns are shifted versions of the hemodynamic response function (HRF) of duration *L* time points at TR temporal resolution, i.e. **h** = [*h*_1_,⋯,*h_L_*]. If *K* echoes are acquired at echo times TE_*k*_, *k* = 1,…,*K*, the signal percentage changes of each echo signal can be vectorized in a column vector of length *NK*. Since the activity-inducing signal can be considered identical for all echoes, the ME signal model can be written as

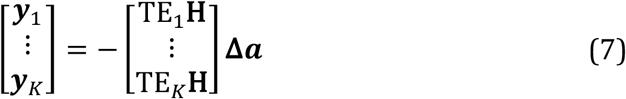

or simply 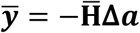.

### Multiecho Sparse Paradigm Free Mapping

The deconvolution algorithm of multiecho sparse paradigm free mapping (ME-SPFM) aims to deconvolve the changes in the BOLD ME-fMRI signal related to neuronal activity without knowledge of their timings. This involves the estimation of **Δ*a*** according to the model in Eq. (7). Figure 1 illustrates a schematic of the assumed ME-fMRI signal model and the ME-SPFM algorithm. Assuming that after preprocessing, the noise follows an uncorrelated Normal distribution, an unbiased estimate of **Δ*a*** can be obtained by means of an ordinary least-squares estimator. Nevertheless, in practice, the least-squares solution would produce estimates with large variability due to the large collinearity between the columns of 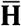. Therefore, it is advisable to incorporate some type of regularization term to the least-squares minimization. Following previous algorithms for the temporal deconvolution of the BOLD fMRI signal, we propose to estimate **Δ*a*** with the following L_1_-norm regularized least-squares estimator

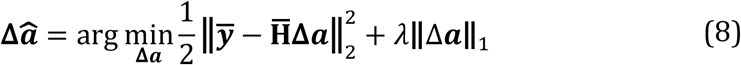

**Figure 1:**
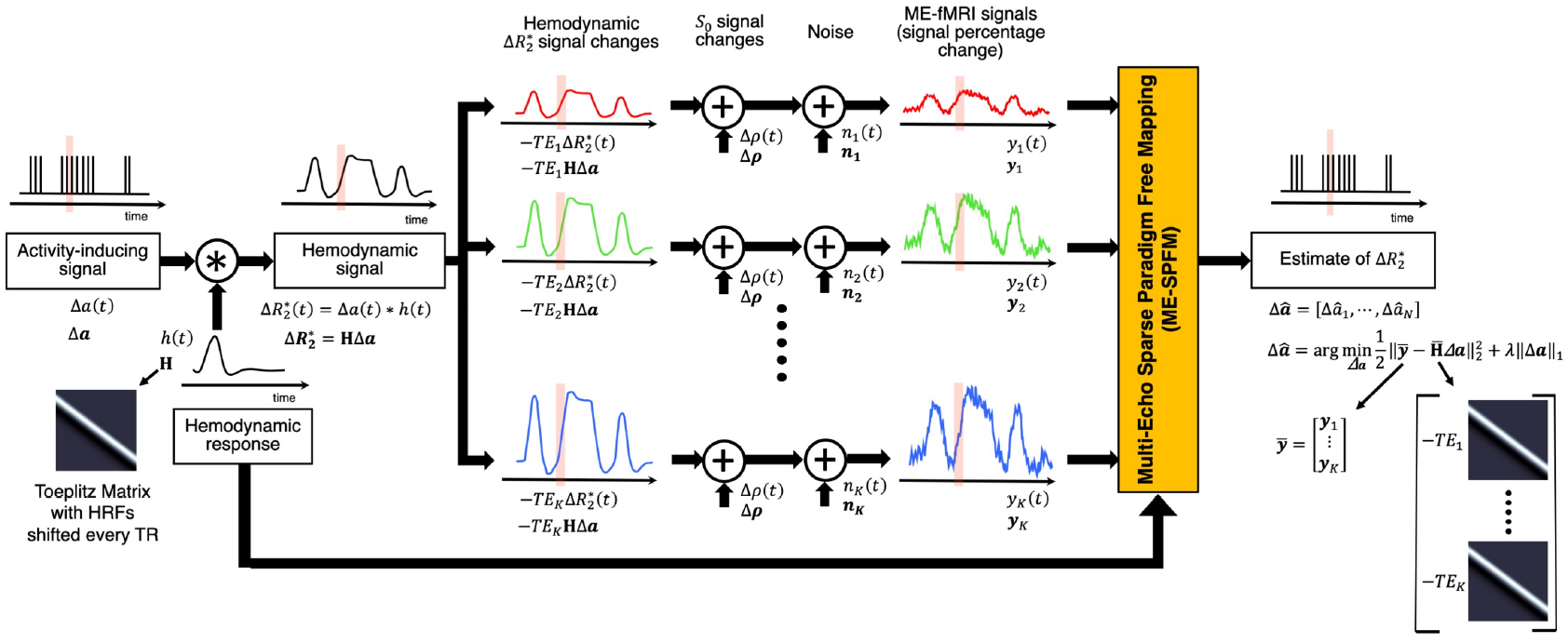
Schematic of the ME-fMRI signal model and the ME-SPFM algorithm. From left to right: An activity-inducing signal (Δa(t)) is convolution with the hemodynamic response (h(t)) resulting in the activity-induced hemodynamic signal or BOLD responses (i.e. 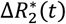). The convolution step can be modelled as multiplying the activity inducing signal with a Toeplitz matrix whose columns are shifted HRFs every TR. Percentage signal changes of the fMRI signal acquired at TE_*k*_ can be modelled as the sum of the hemodynamic signal scaled by TE_k_ (i.e. 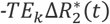), fluctuations of the net magnetization (Δρ(t)), and other noisy sources (e.g. thermal noise) (Δn(t)). The percentage signal changes of all echoes (or their MEICA denoised versions) are concatenated and input to ME-SPFM algorithm, which solves a regularized least squares problem, to obtain estimates of the activity inducing signal.

This mathematical optimization problem is known as Basis pursuit denoising (Chen et al., 1998), which is equivalent to the well-known LASSO (Tibshirani, 1996). The L_1_-norm regularization term encourages sparse estimates with few non-zero coefficients in 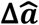, performing both variable selection and regularization in order to enhance the prediction accuracy and the interpretability of the estimates. This implies that 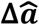 will tend to be non-zero in only the coefficients that explain a large variability of the ME-fMRI signals according to the hemodynamic model.

The choice of the regularization parameter *λ* is critical to obtain an accurate estimate of 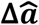. In this work, instead of selecting a fixed value of *λ*, we compute the entire regularization path by means of the least angle regression (LARS) procedure (Efron et al., 2004). This homotopy procedure initializes 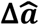 with zero coefficients and then efficiently estimates the entire regularization path for decreasing values of *λ* where a coefficient of 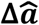 becomes non-zero or shrinks to zero again. After computing the regularization path, we propose to select the estimate of 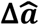 based on the Bayesian Information Criterion (BIC) as follows

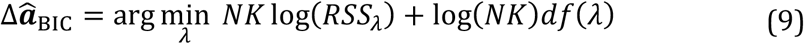

where 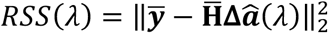 and *df*(*λ*) are the residual sum of squares and effective degrees of freedom for each estimate (Tibshirani and Taylor, 2011; Zou et al., 2007) as a function of *λ*, respectively. Note that the BIC scales with *NK*, i.e. the number of time points by the number of echoes.

Finally, to compensate for the shrinkage towards zero of the coefficients owing to the L_1_-norm regularization term, we propose to perform debiasing of the BIC estimate, known as the relaxed LASSO (Meinshausen, 2007). Debiasing is performed as the ordinary least-squares estimate on the reduced model corresponding to the subset of non-zero coefficients of the estimate. More specifically, let 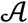 denote the support of 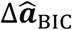, i.e. 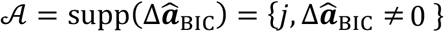, the coefficients of the debiased estimate in the support 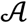 are re-computed as

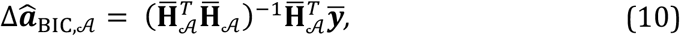

where 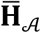 is the reduced matrix with the subset of columns of 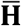 corresponding to the support 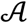, whereas the coefficients not included in 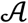 remain as zero.

### MRI data acquisition

The evaluation of ME-SPFM was performed on ME-fMRI data acquired in 10 subjects (5 males, 5 females, mean ± SD age = 25 ± 3 y.o.) using a multi-task rapid event-related paradigm. Six subjects performed two functional runs, and 4 subjects only performed 1 run due to scanning time constraints (i.e. a total of 16 datasets). All participants gave informed consent in compliance with the NIH Combined Neuroscience International Review Board-approved protocol 93-M-1070 in Bethesda, MD. A complete description of the MRI acquisition protocols and experimental tasks in the experimental design can be found in Gonzalez-Castillo et al. (2016), and relevant details are given here for completeness.

MRI data was acquired on a General Electric 3 Tesla 750 MRI scanner with a 32-channel receive-only head coil (General Electric, Waukesha, WI). Functional scans were acquired with a ME gradient-recalled echo-planar imaging (GRE-EPI) sequence (flip angle=70° for 9 subjects, flip angle=60° for 1 subject, TEs=16.3/32.2/48.1 ms, TR=2 s, 30 axial slices, slice thickness=4 mm, in-plane resolution=3×3 mm^2^, FOV 192 mm, acceleration factor 2, number of acquisitions=220). Functional data was acquired with ascending sequential slice acquisitions, except in one subject where the acquisitions were interleaved. In addition, high resolution T1-weighted MPRAGE and proton density images were acquired per subject for anatomical alignment and visualization purposes (176 axial slices, voxel size=1×1×1 mm^3^, image matrix=256×256).

### Experimental paradigm

The PsychoPy software (Peirce, 2009) was used for stimulus delivery. Eye tracking data were collected to check subject’s performance. Each run included 6 trials of each of the 5 different tasks (i.e. a total of 30 trials per run). Subjects were instructed on the task types prior to the scanning session. The 5 tasks were:

1. Finger tapping (FTAP). Subjects were instructed to press one button of a response box with a single finger at a fixed rate of approximately 0.5 Hz for a duration of 4 s. Visual cues were shown to help subjects press the button at a constant rate. All subjects performed this task with the left hand except two, who were inadvertently provided with the response box on their right hand.
2. Biological motion observation (BMOT). Subjects were instructed to observe 4-second videos of dot patterns resembling biological motion such as walking, jumping, dancing, drinking and climbing steps. The videos were shown on only one of the two visual hemi-fields (right or left) and their position was randomized across trials.
3. Passive viewing of houses (HOUS). Subjects were instructed to watch a succession of pictures of houses shown in the center of the screen. Each trial lasted 4 s and contained pictures of 6 different houses. Each house appeared for approximately 170 ms with a gap of 500 ms between pictures.
4. Listening to music (MUSI). Subjects were instructed to attentively listen to 4-seconds recordings of music clips played by a single instrument (violin, piano, or drums) and to direct their gaze to one of the three pictures on the screen (one per instrument) that represented the instrument being played as soon as they had identified it.
5. Sentence reading (READ). Subjects were instructed to covertly read sentences presented on the screen one word at a time. For each trial, words were presented in one of the two hemifields (right or left) to aid with analysis of eye tracking data. All words of a trial appeared on the same hemifield. Each word was presented for 250 ms with gaps of 100 ms in between. Sentence length was between 10 and 11 words, so each trial lasted either 3400 or 3750 ms. Onset times for trials were generated with optseq2 in Freesurfer (https://surfer.nmr.mgh.harvard.edu/optseq). Three different schedules (onset times) were randomly used in these experiments. For all three schedules the minimum inter-stimulus interval (ISI) was 10 s. Mean and standard deviation ISIs for the three different schedules were: 13 ± 24, 13 ± 18 and 13 ± 15 s.

### FMRI data preprocessing

Each ME-fMRI dataset was preprocessed through four different pipelines implemented in AFNI (Cox et al., 1996) resulting in the following datasets:

A. Individually preprocessed echoes (E01, E02 and E03): (1) removal of the initial 10 s to achieve steady-state magnetization, (2) slice timing correction, (3) volume realignment, registration to anatomical image, and warping to MNI template, and computation of the combined spatial transformation, (4) spatial normalization of each echo dataset to the MNI template at 2 mm isotropic voxel size with a single spatial transformation, (5) nuisance regression (Legendre polynomials up to 5^th^ order, realignment parameters and their 1^st^ temporal derivatives, and 5 largest principal components of voxels within the lateral ventricles), (6) spatial smoothing with a 3D Gaussian kernel with Full Width Half Maximum of 6 mm, and (7) calculation of signal percentage change as described in Eq. (4).
B. Optimally combined dataset (OC): same as above, but with optimal weighted combination of the three echoes based on non-linear voxelwise estimation of 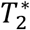 (Posse et al., 1999) between steps (4) and (5).
C. Multiecho Independent Component Analysis plus Optimally Combined dataset (DN): same as E02, but with multiecho independent component analysis (MEICA) denoising (Kundu et al., 2012) and optimal combination between steps (4) and (5). MEICA was applied using the code available in https://github.com/ME-ICA/me-ica (version 3.2).
D. Multiecho Independent Component Analysis denoised echoes (MEICA-E01, MEICA-E02, MEICA-E03): same as above but include MEICA between steps (4) and (5).

### FMRI data analysis

The three preprocessed echo datasets (E01, E02 and E03) and the three MEICA denoised echo datasets (MEICA-E01, MEICA-E02 and MEICA-E03) were analyzed with the ME-SPFM algorithm described above. The ME-SPFM algorithm was implemented for AFNI using functions for compatibility with R and used the LARS package (version 1.2) for the computation of the regularization path of the LASSO. The canonical HRF (SPMG1 option of 3dDeconvolve in AFNI) was used as the hemodynamic response function (HRF) to define the convolution matrices in 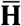. Since the ME-SPFM algorithm outputs a 4D-dataset with voxelwise time-varying estimates of **Δ*a***, which are related to 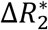, for validation purposes we defined ME-SPFM activation maps for each trial by computing the maximum of the **Δ*a*** volumes when each trial occurred (i.e. 3 TRs for a duration of 4 s per trial).

The performance of ME-SPFM was compared with the results of the deconvolution with the Sparse Paradigm Free Mapping (1E-SPFM, Caballero-Gaudes et al., 2013) and traditional GLM analyses implemented with 3dREMLfit in AFNI. These analyses were performed in the E02, OC and DN datasets. As for 1E-SPFM, datasets were analyzed with the implementation of SPFM available in AFNI (3dPFM program) using the LASSO algorithm, the Bayesian Information Criterion (BIC) for selection of the regularization parameter and the canonical HRF to define the corresponding convolution matrix. Similar to ME-SPFM, 1E-SPFM activation maps were created from the deconvolved coefficients (beta output dataset in 3dPFM) as the maximum of the volumes when each trial occurred. No additional processing steps were applied to the ME-SPFM and 1E-SPFM activation maps.

In addition, we performed two different GLM analyses in the E02, OC and DN datasets, where the design matrix was either defined considering all trials of a task in one regressor (TASK-LEVEL) or each trial individually modulated, i.e. each trial has its own regressor (‘IM’ or TRIAL-LEVEL). The SPM canonical HRF was used in both analyses, assuming a trial duration of 4 s. The task-based activation maps were thresholded at FDR-corrected *q* ≤ 0.05 (TASK-q05). The trial-based activation maps were thresholded at FDR-corrected *q* ≤ 0.05 (IM-q05), as well as uncorrected *p* ≤ 0.05 (IM-p05) and *p* ≤ 0.001 (IM-p001). The number of components removed by MEICA was considered in the computation of the degrees of freedom of the GLM analyses of the DN dataset.

### Evaluation of spatial concordance with GLM analyses

We evaluated the ability of the 1E-SPFM and ME-SPFM to detect the activation revealed by the GLM analyses in terms of the spatial sensitivity, spatial specificity and spatial overlap using a dice coefficient metric. This evaluation only considered activations that produce a positive BOLD signal change (i.e. a positive effect size in GLM analyses), a positive coefficient in 1E-SPFM and a negative 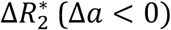 coefficient in ME-SPFM.

First, we performed the comparison at the task level by using the task-based activation maps obtained with the DN dataset (TASK-q05/DN) as the reference maps. These can be considered as the gold standard of activation maps per task in each dataset that can be obtained with an analysis that is aware of the trials’ onsets and durations using the same hemodynamic model (i.e. SPMG1). For this comparison, we also considered the following activation maps: IM-q05, IM-p001, IM-p05 and 1E-SPFM for E02, OC and DN input datasets, as well as ME-SPFM using the triplets E01, E02 and E03 or MEICA-E01, MEICA-E02 and MEICA-E03 as input datasets.

Second, we performed the comparison at the trial-level by using the trial-based activation maps at *p* ≤ 0.05 obtained with the DN dataset (IM-p05/DN) as the reference maps, which considers a model of a single trial based on its onset and duration and, thus, is closer to the assumptions of the ME-SPFM activation maps. For this comparison, we considered the following activation maps: 1E-SPFM for E02, OC and DN datasets, as well as ME-SPFM using the triplets E01, E02 and E03 or MEICA-E01, MEICA-E02 and MEICA-E03 as input datasets.

### Evaluation of temporal concordance with GLM-IM analysis

We also computed maps of the Pearson correlation between the fitted signal of the GLM-IM model with the fitted signal of 1E-SPFM and ME-SPFM (i.e. convolution of the detected events with the canonical HRF) in order to evaluate the temporal concordance of the detected events in comparison with a conventional model-based analysis using timing of the experimental trials. Moreover, we computed the correlation between these models with the preprocessed DN (i.e. MEICA+OC) dataset to examine whether the deconvolution approaches can explain additional variance of the preprocessed data, particularly in regions that might not be involved during the known tasks. These temporal correlation analyses can serve as an evaluation criterion that is not threshold-dependent and therefore complements the aforementioned spatial evaluation.

### Quantitative Analysis of 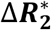 estimates

We evaluated the ability of ME-SPFM deconvolution to estimate 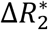 changes that range within physiologically plausible limits, which was established as 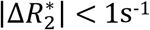 according to previous reports of neurobiologically-driven 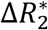 values at 3T (van der Zwaag et al., 2009). First, we computed histograms of 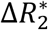 estimates for both ME-SPFM and MEICA-ME-SPFM activation maps in three conditions: a) during the entire dataset in all whole-brain voxels to assess the efficacy of the algorithms to yield physiologically-plausible estimates independently of the paradigm, b) during the timings of trials in all whole-brain voxels to examine whether both positive and negative 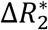-estimates occur with each trial, and c) during the timings of trials in only those voxels showing positive activation in the TASK-q05 maps for the DN dataset, i.e. assumed to have a clear positive BOLD response to the task that is associated with 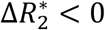. In addition, we computed the percentage of estimates exceeding 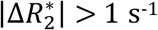 according to these three options per trial, per task, per dataset and per ME-SPFM analysis.

## RESULTS

The output of ME-SPFM is a 4D dataset with an identical number of time points as the input dataset, which can be visualized as a sequence of deconvolved maps. The movies showing the 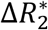 and 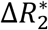-fitted signals for all the runs and subjects are available in https://ccaballero.pages.bcbl.eu/me-spfm-videos/. Figure 2 depicts the corresponding activation maps for representative single trial events of each task type for the same run for IM-p05, 1E-SPFM with the DN dataset, and ME-SPFM with the preprocessed echo datasets and after ME-ICA. The 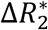-activation maps obtained with ME-SPFM have a larger resemblance with the maps obtained with the trial-based GLM analysis (IM-p05) than the activations maps obtained with 1E-SPFM. Even though the 1E-SPFM maps generally depict clusters of activation in the same locations as the GLM maps (i.e. high spatial specificity), they exhibit lower spatial sensitivity than the ME-SPFM activation maps, especially observed in FTAP-5, HOUS-1 and READ-2. In general, the ME-SPFM activation maps exhibit negative 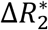 in the brain regions showing positive signal changes, and conversely positive 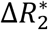 in brain regions showing negative signal changes, in the IMp05 and 1E-SPFM maps. The MEICA+ME-SPFM activation maps illustrate that applying MEICA prior to ME-SPFM reduces spurious activations in the borders of clusters and draining veins, probably related to inflow fluctuations, and in brain edge voxels related to effects of head motion. Some examples of these effects are marked with green arrows.

**Figure 2:**
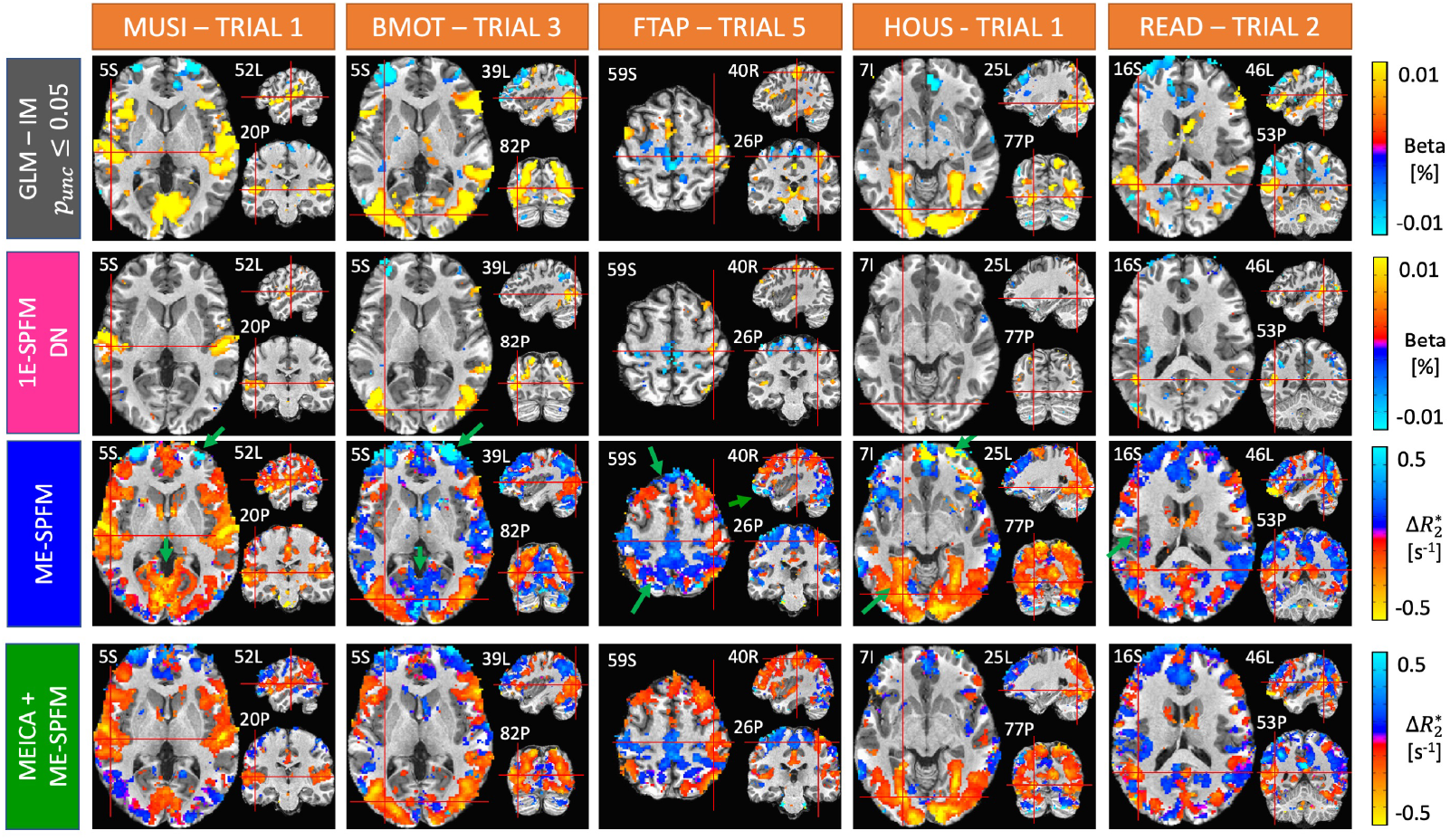
Activation maps of an individual single-trial event for each experimental condition obtained with an individually-modulated (IM) GLM analysis (T-test, uncorrected p ≤ 0.05) (first row), 1E-SPFM (second row), ME-SPFM (third row) and MEICA+ME-SPFM (fourth row). The maps of IM-GLM and 1E-SPFM show estimated beta coefficients in signal percentage change (i.e. % amplitude), whereas ME-SPFM and MEICA-ME-SPFM show estimated 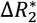 values in units of s^−1^. Note the ME-SPFM and MEICA+ME-SPFM activations maps are shown with reverse colorbars so that negative (positive) 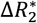 values shown in red (blue) induce positive (negative) BOLD signal changes.

Figure 3 plots the preprocessed and estimated signals for the trial-based GLM and MEICA+ME-SPFM analyses for the same dataset in seven representative voxels relevant to each task (shown in the left maps), as well as in left precuneus and right dorsolateral prefrontal cortex (DLPFC), which are regions typically associated with the default mode network and dorsal attention network, respectively. For the voxels related to the tasks, the 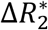 time series obtained with ME-SPFM exhibit negative 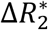 (deflection in blue traces) that are reliably detected at the time of all experimental events (light green bands). These negative 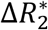 result in positive BOLD signal changes (red traces) that agree with the signals estimated by the GLM-IM model in the DN dataset (green traces). The Pearson’s correlation between the BOLD signal estimated with ME-SPFM and with the GLM-IM model is shown on top of each time course (Corr-FIT values). Clearly this correlation is larger in the task-activated voxels than in the left precuneus and right DLPFC. In addition to the task-related events, ME-SPFM is able to detect BOLD events that often occur during the timing of other tasks or in the absence of any task, i.e. during rest periods. The 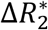 activation maps shown on the right correspond to five representative task-unrelated spontaneous BOLD events marked with red dashed lines in the plots of the left precuneus and DLPFC. These maps depict spatial patterns with clusters of activation in regions of the default mode network (negative 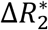 at 58, 132, 316 and 404 s and positive 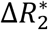 at 92 s) that act in synchrony with clusters of activation in areas of the dorsal attention network. Importantly, these transient events occurring at times without any task cannot be revealed by conventional GLM approaches, but are detected by ME-SPFM owing to its ability to operate without timing information.

**Figure 3:**
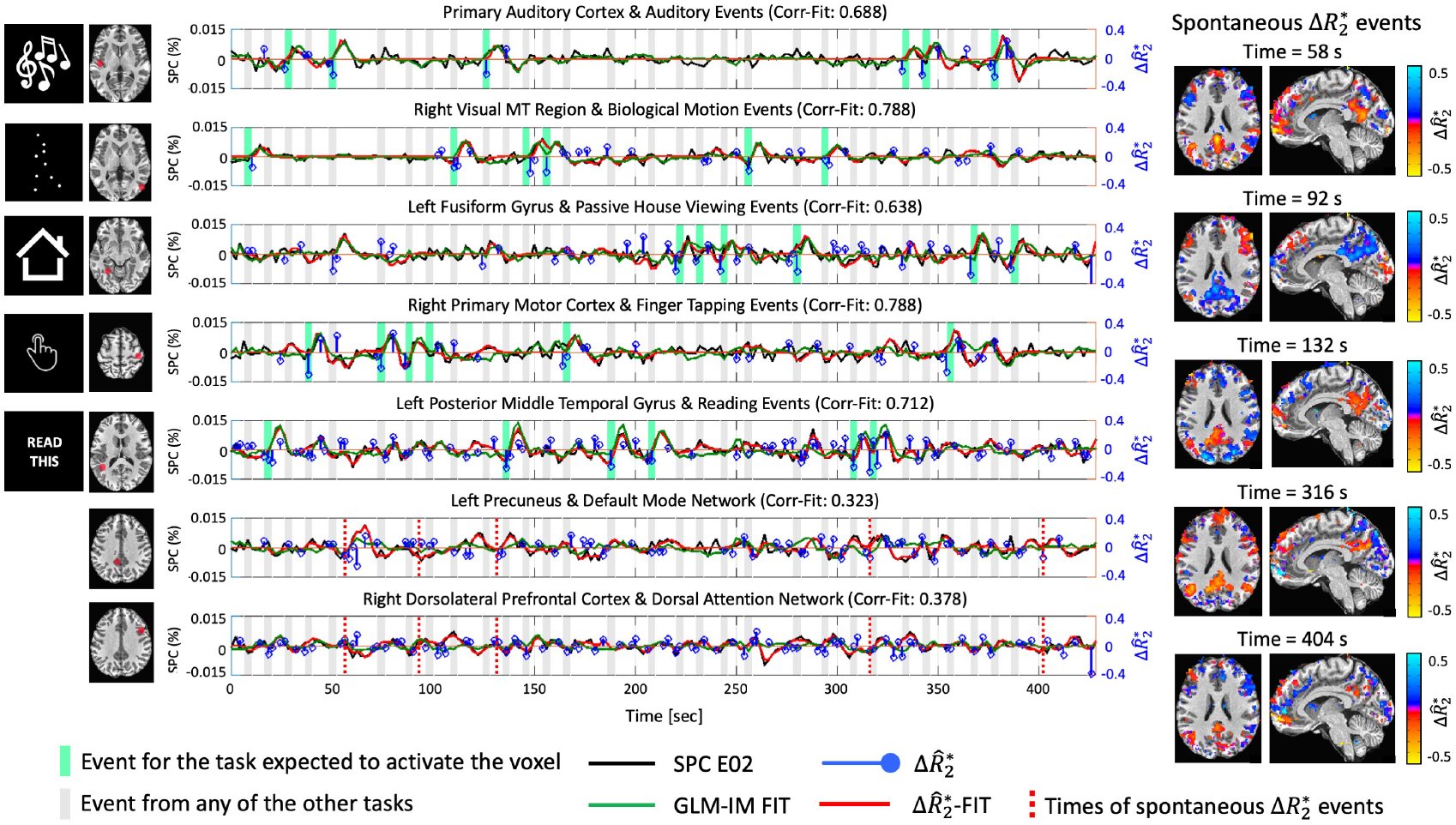
Time courses of signal percentage change in E02 dataset (SPC E02, black line), GLM-IM fitted signal (green), 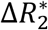 (blue) and 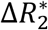 BOLD estimates, i.e. 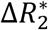 convolved with canonical HRF (red) obtained with MEICA+ME-SPFM in seven representative voxels in task-related regions, and left precuneus and right dorsolateral prefrontal cortex (DLPFC) of the same dataset as Figure 1. The voxel’s location is shown in the left maps. Dark and light grey bands indicate the times of trials of the relevant task for each voxel and the rest of the tasks, respectively. The maps shown on the right display instances of spontaneous 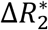 events occurring at rest, whose timing is marked with dashed lines in the time courses of the left precuneus and right DLPFC.

Considering all datasets, Figures 4 and 5 show the average spatial dice coefficient, sensitivity and specificity for the different methods using the task-based GLM (TASK-q05/DN) and trial-based GLM maps (IM-p05/DN) as reference maps, respectively. Both figures illustrate the ME-SPFM algorithm outperforms its 1E-SPFM counterpart regardless of the prior use of ME-ICA, achieving considerably larger spatial overlap and sensitivity with a reduction in specificity. As shown in Figure 4, ME-SPFM achieves similar spatial concordance with the TASK-q05 maps to the one obtained with trial-based GLM analyses when the statistical significance threshold is set between *p_unc_* ≤ 0.05 (IM-p05) and *p_unc_* ≤ 0.001 (IM-p001). In general, denoising the fMRI signal with MEICA (DN dataset) is beneficial to increase the sensitivity and the spatial concordance of individual GLM TRIAL-LEVEL maps with respect to the TASK-LEVEL maps. In all cases, the spatial concordance of the IM-p001 maps is similar to the TRIAL-q05 maps. MEICA-based denoising is more advantageous than preprocessing based on optimal combination of echoes (OC dataset) or single-echo (E02 dataset) for detecting single-trial BOLD events in both 1E-SPFM and ME-SPFM analyses. The advangage of MEICA is also seen in the IM-p05 maps. Similar conclusions can be drawn from the results in Figure 5 wherein the TRIAL-LEVEL IM-p05/DN activation maps become the reference maps. MEICA+ME-SPFM yields larger spatial concordance, sensitivity and specificity than ME-SPFM, and both of them outperform 1E-SPFM analyses in terms of spatial overlap for all the conditions.

**Figure 4:**
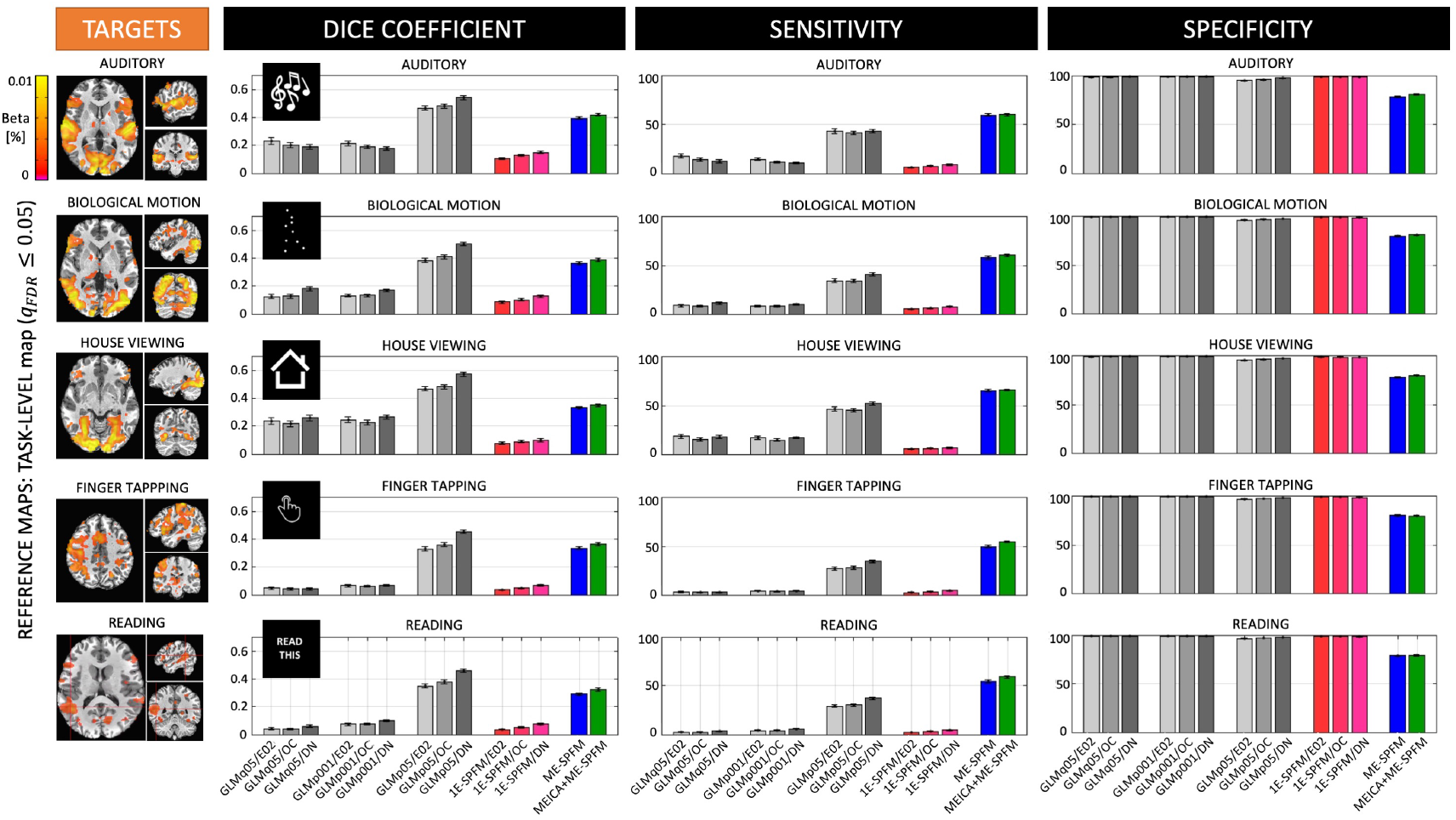
Average values of dice coefficient (i.e. spatial overlap), sensitivity and specificity across all the singletrial activation maps obtained with each of the analysis methods (GLM-IM, 1E-SPFM, ME-SPFM and MEICA+ME-SPFM) for each of the experimental conditions. TASK-LEVEL activation maps thresholded at FDR-corrected q ≤ 0.05 (TASKq05) and only including voxels with positive activation were used as reference maps, shown on the left for a representative dataset. The GLM-IM and 1E-SPFM activation maps were computed from the E02, OC and DN (i.e. MEICA and OC) preprocessed datasets, and GLM-IM maps were obtained at thresholds FDR-corrected q ≤ 0.05 (IMq05), as well as uncorrectedp ≤ 0.05 (IMp05) and p ≤ 0.001 (IMp001).

**Figure 5:**
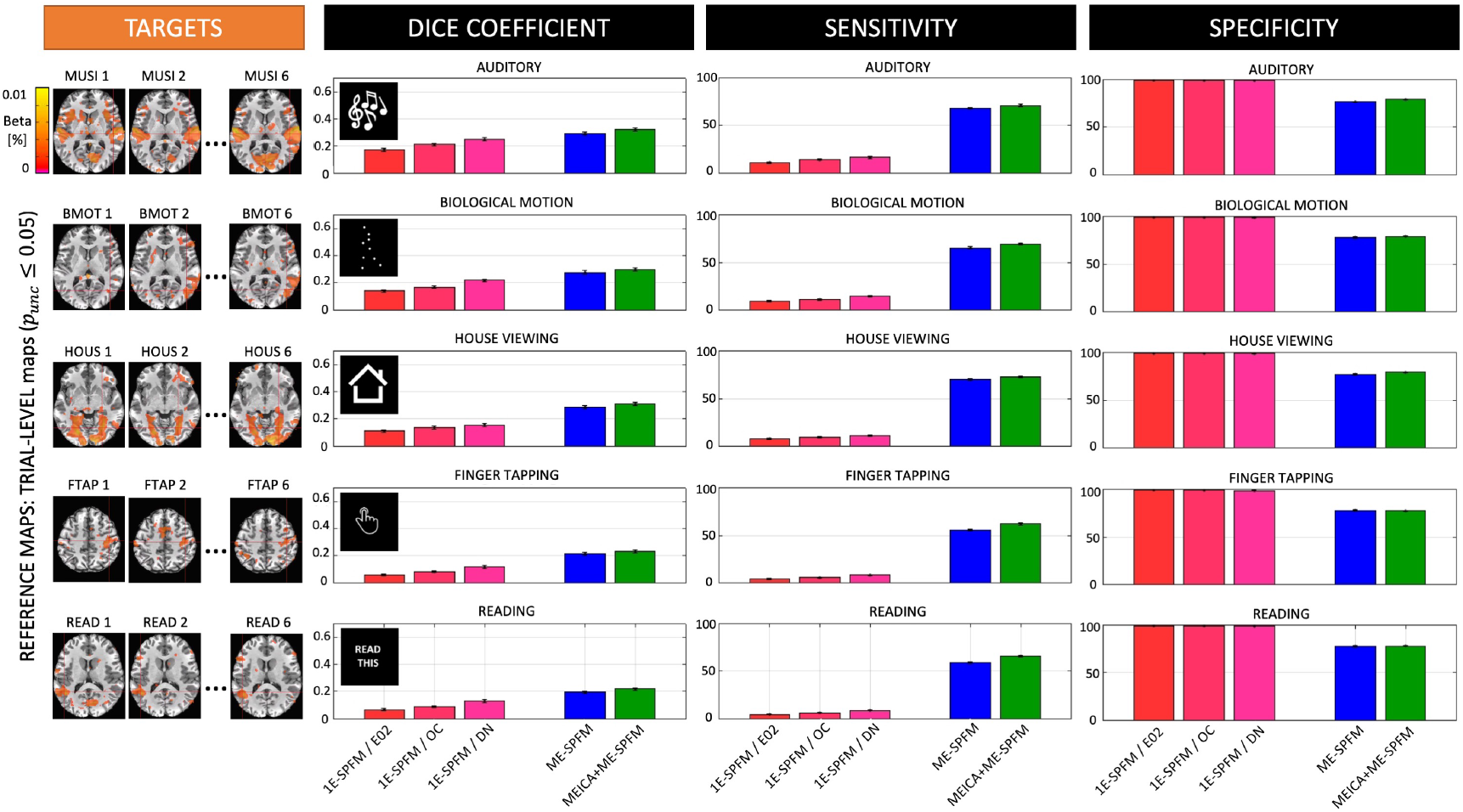
Average values of dice coefficient (i.e. spatial overlap), sensitivity and specificity across all the singletrial activation maps obtained with 1E-SPFM, ME-SPFM and MEICA+ME-SPFM for each of the experimental conditions. TRIAL-LEVEL activation maps thresholded at uncorrected p ≤ 0.05 (IMp05) and only including voxels with positive activation were used as reference maps, shown on the left for a representative dataset. The 1E-SPFM activation maps were computed from the E02, OC and DN (i.e. MEICA and OC) preprocessed datasets.

Figure 6 includes the receiver operating characteristic (ROC) curves with the sensitivity and specificity of each individual trial’s activation map for all conditions and the two types of reference maps: TASK-q05/DN are shown at the top and IM-p05/DN are shown at the bottom. For visualization purposes, only the IM-q05, IM-p001 and IM-p05 with the DN dataset are shown in the ROC plots at the TASK-LEVEL. The radius of each circle is relative to the number of voxels showing activations (i.e. total number of positives) in the reference maps. Similar to Figures 4 and 5, the ROC curves illustrate that ME-SPFM offers larger sensitivity in detecting single-trial events than 1E-SPFM, which instead achieves nearly perfect specificity values (i.e. above 95%) similar to GLM-IM activation maps. Interestingly, ME-SPFM achieves larger sensitivity values than GLM-IM for certain trials, particularly for the house viewing and reading conditions. The use of MEICA in preprocessing slightly improves the performance of ME-SPFM, particularly when compared with the TRIAL-LEVEL IM-p05/DN activation maps.

**Figure 6:**
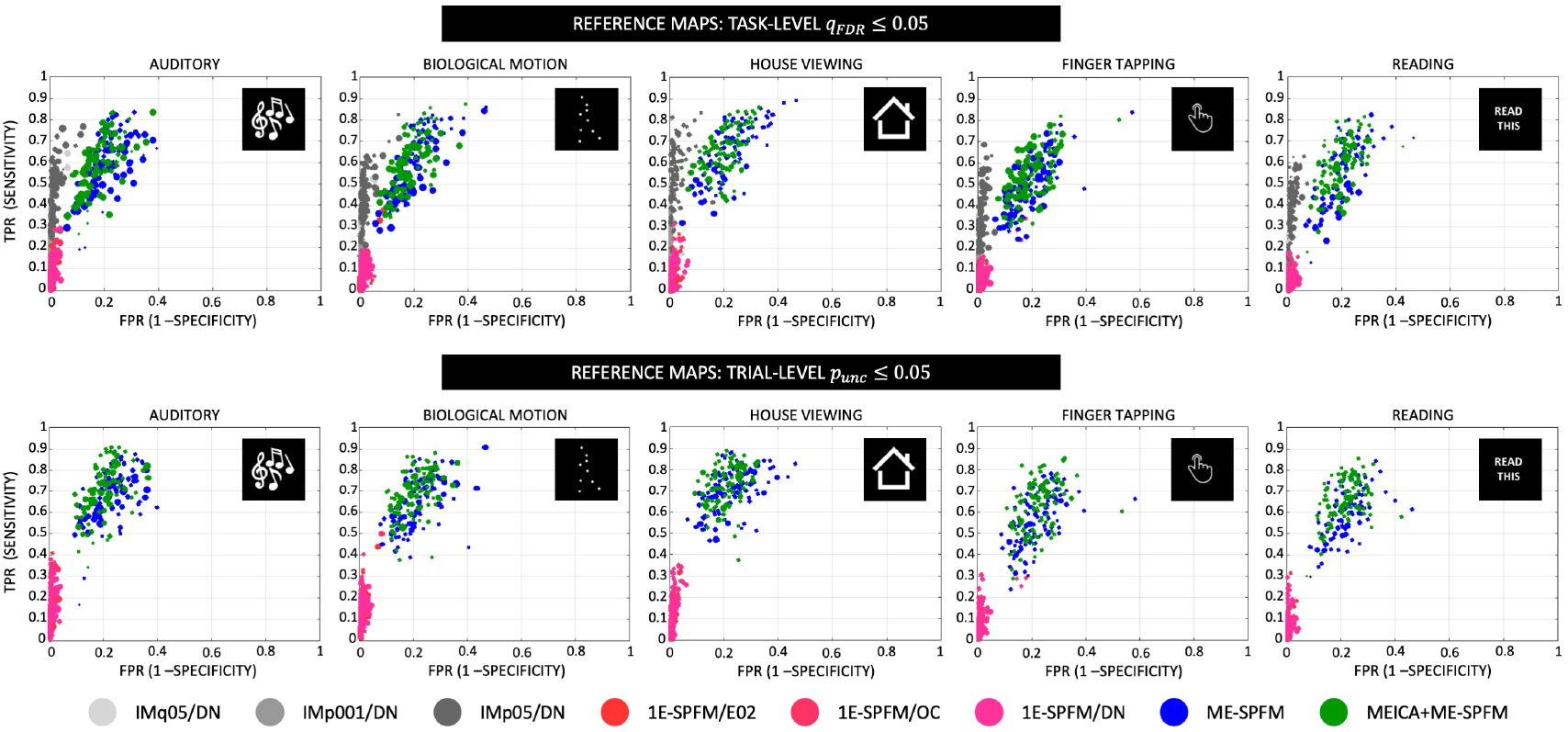
Receiver operating characteristic (ROC) curves with the sensitivity and specificity of each individual trial’s activation map for all conditions and the two types of reference maps: TASK-q05/DN (top) and IM-p05/DN (bottom). For visualization purposes, only the IM-q05, IM-p001 and IM-p05 with the DN dataset are shown in the ROC plots at the TASK-LEVEL. The radius of each circle is relative to the number of positives in the reference map, i.e. trials with bigger circles activations had more activated voxels in the reference maps, wherein the largest radius is the maximum number of positives across all trials and conditions.

To demonstrate the temporal concordance of the detected activation, Figure 7 shows the average Pearson’s correlation coefficients of the fitted signal estimated with the GLM-IM model (top three rows) and the preprocessed DN dataset (bottom four rows) with the fitted signal obtained with 1E-SPFM/DN, ME-SPFM and MEICA+ME-SPFM. Fisher’s z-transformation was applied to the correlation coefficients prior to averaging across datasets and then inversely applied for visualization purposes. Also, notice that the range of the correlation maps only covers positive values because negative correlations were only identified in few disperse voxels in white matter. As for the correlation with the GLM-IM fitted signal, both ME-SPFM analyses show higher temporal correlation values than those obtained with 1E-SPFM, particularly confined to gray matter voxels. The peaks of the correlation maps occur in brain regions involved in the processing of the multiple tasks, such as the primary auditory cortex for listening to music, the primary motor cortex for finger tapping, the ventral occipitotemporal cortex involved for viewing of houses and reading, the posterior temporal-occipital cortex for passive viewing of biological motion, and the primary occipital cortex for the multiple tasks with visual input. The 1E-SPFM maps only display large correlation values in these cortical regions.

**Figure 7:**
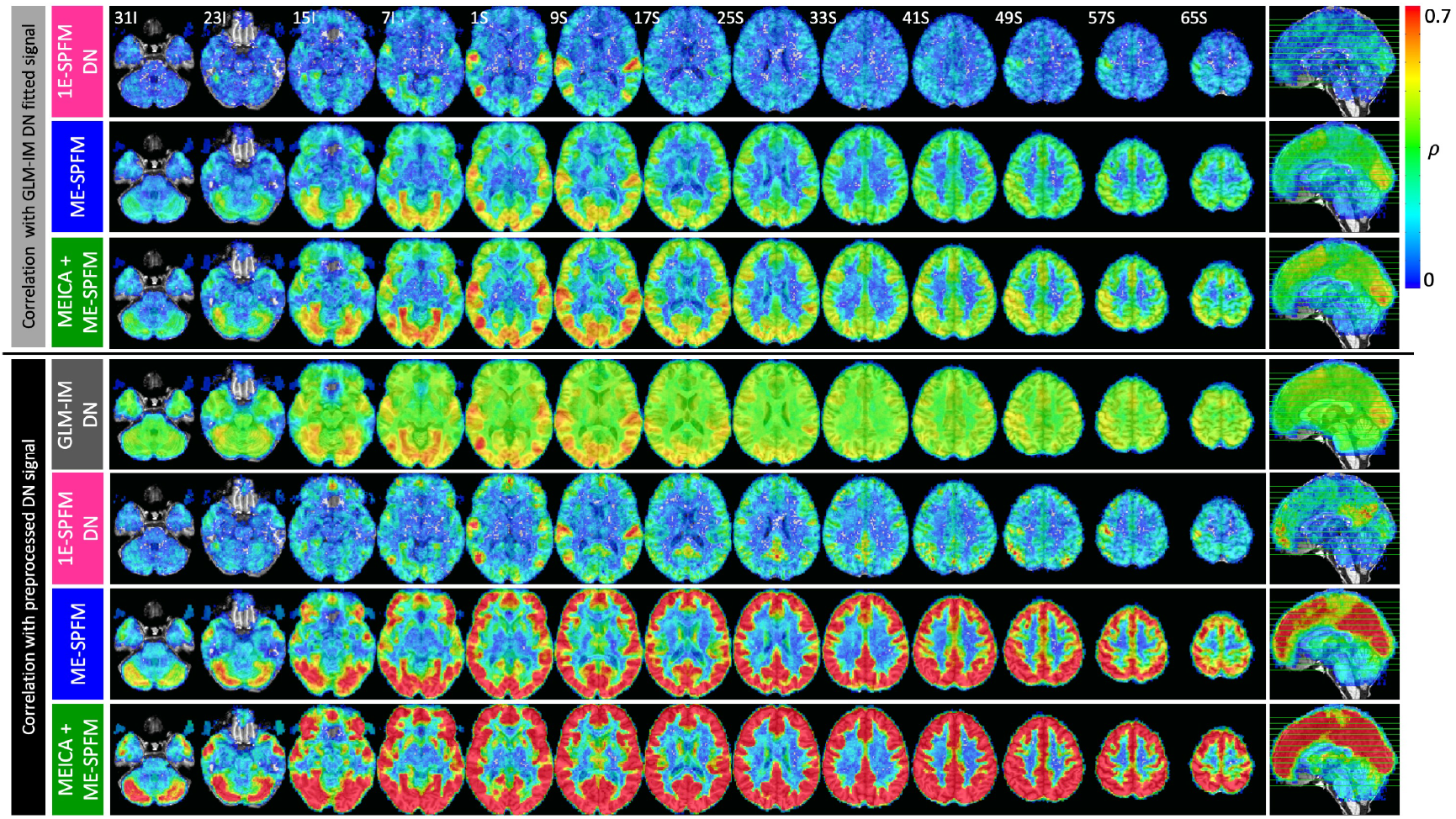
(Top three rows) Maps of Pearson’s correlation coefficients between the BOLD signals estimated with the GLM-IM analysis and the BOLD signals estimated with the 1E-SPFM, and ME-SPFM and MEICA+ME-SPFM deconvolution algorithms. (Bottom four rows) Maps of Pearson’s correlation coefficients between the preprocessed DN dataset and the fitted signals estimated with the GLM-IM analysis, and the 1E-SPFM, and ME-SPFM and MEICA+ME-SPFM deconvolution algorithms.

The bottom four rows of Figure 7 illustrate the average correlation maps of the fitted signals with the preprocessed DN dataset. The correlation maps of the DN dataset with the GLM-IM fitted signal are spatially smooth with non-negligible correlation across all brain voxels and peaks in task-related areas. Similar to the correlation maps with the GLM-IM fitted signals, the ME-SPFM correlation maps reveal a pattern of larger correlation values in voxels across the entire cortex and in some subcortical areas (e.g. putamen and caudate nucleus) and low correlation values in voxels in white matter and cerebrospinal fluid where the deconvolution normally produces null estimates. The correlation values of the MEICA+ME-SPFM fitted signal with the DN dataset is larger than those obtained with the ME-SPFM maps. This can be expected since the reference signal has also been denoised with MEICA. Both ME-SPFM clearly exhibit larger correlations than the 1E-SPFM due to their higher temporal sensitivity. The widespread pattern of correlation in grey matter can also be observed, but less evidently so in the maps obtained with 1E-SPFM. The deconvolution approaches are able to explain variance of the preprocessed DN signal in regions such as the anterior and posterior cingulate cortices, precuneus and prefrontal regions, which cannot be described with the GLM-IM model.

Figure 8 displays the histograms of 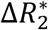 values estimated with ME-SPFM and MEICA+ME-SPFM for all subjects: (A) during the entire run, (B-F) during the timings of each task in all intracranial voxels, and (G-K) only in the voxels with significant positive response in the corresponding TASK-q05/DN activation map. Voxels with zero 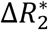 are discarded in the histogram plots. In general, 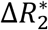 estimates fall within the range of [−1, 1] s^−1^, which is a physiologically-plausible range of 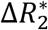 in grey matter at 3T. In addition, the percentage of voxels with 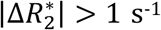 was considerably reduced in the MEICA+ME-SPFM analyses (see plots L and M). A table with the percentage of voxels with 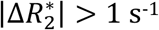 for ME-SPFM and MEICA+ME-SPFM for all datasets is available as supplementary material. The histograms illustrate that the MEICA+ME-SPFM 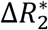-estimates have smaller amplitude than the ME-SPFM 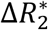-estimates. Furthermore, the histograms become skewed towards negative 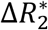-estimates when the mask only includes voxels with significant positive task-related BOLD signal changes. Interestingly, the histograms exhibit a noticeable symmetry around 0 s^−1^ with all intracranial voxels, particularly when the entire duration of the run is considered.

**Figure 8:**
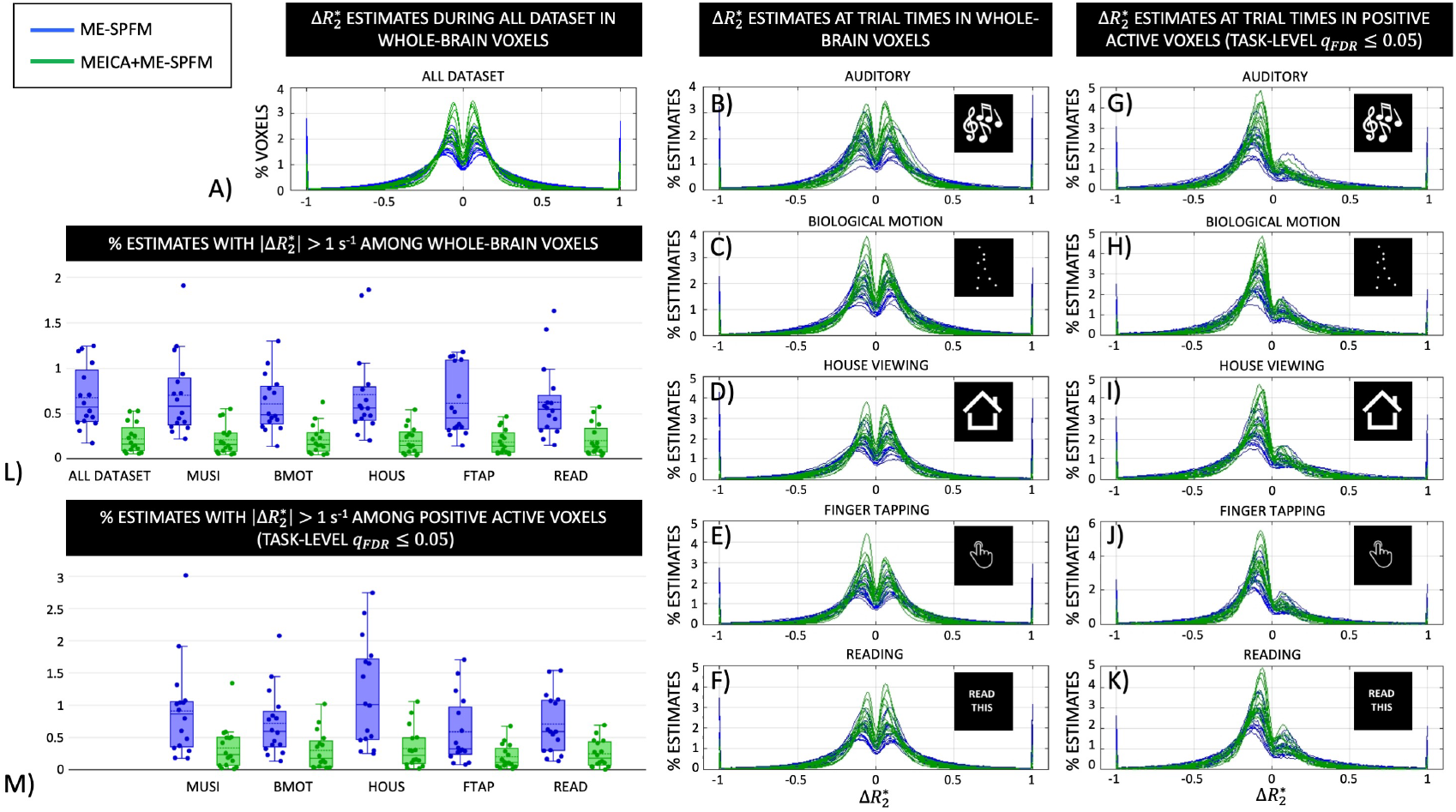
Histograms 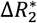 values estimated with ME-SPFM (blue lines and boxes) and MEICA+ME-SPFM (green lines and boxes) in: A) whole-brain voxels during the entire dataset, (B-F) in whole-brain voxels during times of trials for each condition, and (G-K) in voxels with positive activation according to the TASK-LEVEL IMq05 activation map during the times of trials for each condition. (L-M) Box plots with the percentage of voxels with 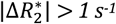 for the analysis, showing one circle per dataset.

## DISCUSSION

The proposed deconvolution algorithm for ME-fMRI, named multi-echo sparse paradigm free mapping (ME-SPFM), achieved larger spatial overlap with a map obtained using GLM and greater sensitivity than single echo deconvolution but reduced specificity relative to its 1E-SPFM counterpart (Caballero-Gaudes et al., 2013). Even though the deconvolution with 1E-SPFM generated single-trial activation maps with very high specificity, it exhibited a significant reduction in sensitivity that caused the algorithm to fail in the detection of activations in brain regions related to the task for certain events (see 1E-SPFM activation map of HOUS Trail 1 in Figure 2) probably due to insufficient contrast-to-noise ratio in these trials. Here, the deconvolution with ME-SPFM was performed with the same combination of sparsity-promoting regularized estimator of LASSO and the Bayesian Information Criterion as for 1E-SPFM. Thus, it can be inferred that the superior performance of ME-SPFM is due to its ME-based formulation as this accounts for the linear dependence of the BOLD signal on TE according to a monoexponential decay model. Importantly, the advantage of ME-SPFM over 1E-SPFM was observed for the three ways of preprocessing to generate a single dataset from the multiple echo datasets, even after MEICA denoising and optimal echo combination, which can be considered one of the most advanced preprocessing approaches for ME-fMRI data (Gonzalez-Castillo et al., 2016). It can be concluded that, for the purpose of voxelwise deconvolution, leveraging the information available across the multiple echo datasets through a TE-dependent model is more advantageous than MEICA denoising (Kundu et al., 2012) or weighted combination of the multiple echoes in a single dataset (Posse et al., 1999; Gowland and Bowtell, 2007; Poser et al., 2006). In addition, the advantage of ME-SPFM with respect to 1E-SPFM increased when the reference activation maps were defined based on a GLM analysis with task-level regressors, rather than with individually modulated regressors for each trial. This result may be explained due to the higher level of uncertainty of the single-trial activation maps used as reference, which results in higher variability in the specificity, sensitivity and spatial overlap estimates.

We observed that applying MEICA prior to ME-SPFM did improve, but not substantially, the sensitivity, specificity and spatial overlap with the TASK-LEVEL or TRIAL-LEVEL reference maps. Hence, we conclude that the improved ability to blindly detect individual BOLD events is more associated with the ME-SPFM algorithm rather than due to denoising with MEICA denoising. To some degree, this result also demonstrates that the proposed ME-SPFM algorithm can cope with *S*_0_-related fluctuations of the signal despite these being neglected in the deconvolution. Moreover, the slight improvements in performance of the 1E-SPFM and ME-SPFM algorithms when the echo datasets are denoised with MEICA are similar to the ones observed in the GLM analyses, which is concordant with previous results (Gonzalez-Castillo et al., 2016).

Denoising the echo datasets with MEICA prior to the proposed ME-SPFM algorithm is still recommended (i.e. the MEICA+ME-SPFM analysis), since the corresponding activation maps become more focal, showing a reduced number of voxels with non-zero 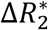 values that may originate from inflow effects, movement-related artefacts and physiological fluctuations (see green arrows in Figures 1). Moreover, the number of voxels with non-physiologically plausible 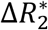-estimates is significantly reduced in MEICA+ME-SPFM (see Figure 8). The reduction in amplitude of the MEICA+ME-SPFM 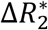-estimates also agrees with previous observations of diminished effects sizes in task-related activations observed in Gonzalez-Castillo et al. (2016).

As shown in Figure 7, the ME-SPFM algorithm also exhibited higher temporal correlation with the GLM-IM fitted signals than 1E-SPFM, suggesting a higher temporal sensitivity of the estimated 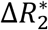. Even though the peaks of the temporal correlation maps were located in brain regions assumed to strongly engage in the experimental tasks, the correlation maps of ME-SPFM also exhibited non-negligible correlation values in grey matter voxels across the entire cortex, subcortical regions and cerebellum, whereas the correlation was clearly reduced in white matter voxels. This indicates that ME-SPFM offers not only higher temporal sensitivity, but also does not detect 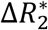-events at random that are specific to brain regions of potential functional relevance. These 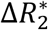-events are missed by 1E-SPFM and cannot be explained from the experimental design with GLM analyses. There can be multiple causes for the origin of these activations. First, the higher contrast-to-noise ratio of the BOLD signal in grey matter voxels than in white matter (Krüger and Glover, 2001). Second, due to the sluggishness of the hemodynamic response, BOLD signal changes associated with 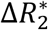 occurring prior to the trials may also extend in time and overlap with the BOLD signal changes in response to the trials. Third, part of the activations observed in brain regions beyond those primarily involved in the performance of the tasks could also be explained in terms of behavioural differences across trials, for instance due to changes in attention, self-awareness or executive control mechanisms that engage other brain regions in a less prominent manner and thus are only detected with ME-SPFM due its enhanced sensitivity relative to 1E-SPFM. Here, we confirmed that subjects performed all task-related events based on eye-tracking measurements, thus ensuring that variability across trials is not associated to inappropriate performance of the tasks (Gonzalez-Castillo et al., 2016).

Importantly, ME-SPFM also enabled us to detect 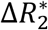-events in periods between trials when the subjects are not assumed to engage in any evoked task. These spontaneous events detected during rest would be neglected by any analysis approach that only model events with timing known by the experimenter. Figure 3 shows several instances of these transient, spontaneous 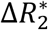-events for a representative dataset in brain regions of the default mode network (Raichle, 2015) as well as the attention and frontoparietal executive control networks (Dixon et al., 2018; Fox et al., 2006). Similar patterns of spontaneous 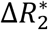-events were observed across all datasets. The maps and amplitude of these spontaneous activations highly resemble the functional connectivity maps observed in resting state fMRI and also exhibit similar between-network relationships in the sign of the detected activations. For instance, these illustrative maps show the well-known opposite polarity of BOLD signal changes, and thus also in 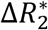, between regions of the default mode network (i.e. precuneus, posterior cingulate, inferior parietal lobule and medial prefrontal cortex) and the dorsal attention network (i.e. dorsolateral prefrontal cortex, frontal eye fields, intraparietal sulcus, superior parietal lobule) (Fox et al., 2005). Although the datasets were not acquired in resting state, these findings corroborate previous evidence of involvement of resting state functional networks during event-related paradigms obtained in single-echo datasets with SPFM (Caballero-Gaudes et al., 2013; Petridou et al., 2013) and Total Activation (Karahanoğlu et al., 2013). Point process analyses have also revealed the presence and relevance of these extreme events in resting-state analyses (Liu et al., 2018; Tagliazucchi et al., 2012; Tagliazucchi et al., 2016).

The 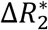 signals estimated with ME-SPFM have interpretable units in s^−1^. As shown in Figure 8, most of the ME-SPFM estimates fell within limits of neuronally-driven 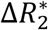 at 3T. For positive BOLD signal changes, Donahue et al. (2011) and van der Zwaag et al. (2009) reported total 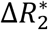 values of −0.74 ± 0.05 s^−1^ and −0.98 ± 0.08 s^−1^ in the human visual and motor cortices at 3T for a block-design tasks with long stimuli. These values are higher than those obtained with our deconvolution in response to more complex tasks and with shorter trial duration. Should we have a prior hypothesis of the maximum 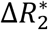 induced by neuronally-driven events per brain region, this information can be exploited to characterize the nature of the detected events and identify those events with exceeding 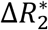 values that might be more related to artefactual changes in the BOLD signal than to neurobiological processes. For that, it is important to consider that 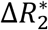 values may vary due to differences in anatomy across brain regions (e.g. vascularization), imaging parameters (e.g. magnetic field strength, RF coil type, voxel size, flip angle) and experimental paradigm (e.g. block vs. fast event-related designs).

Furthermore, the histograms of 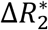-estimates were symmetrical at the whole-brain level, particularly when considering the entire run. The symmetry remained when a gamma function (GAM option in 3dDeconvolve) without post stimulus undershoot was used for deconvolution (data not shown), indicating that it cannot be completely explained due to spurious estimates that try to compensate mismatches between the assumed and real HRF shapes. This hemodynamic equilibrium in number and magnitude is intriguing, but agrees with our previous observations of whole-brain, widespread activations with equivalent number of positive and negative BOLD signal changes at 3T (Gonzalez-Castillo et al., 2012). Although this work does not aim to explore this issue, we conjecture that the observed hemodynamic equilibrium has a main neuronal contribution probably due to inhibition (Devor et al., 2007; Shmuel et al., 2006), rather than purely hemodynamic due to blood stealth of positively active regions from neighboring regions (Harel et al., 2002). Our support to this claim is that positive 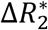-estimates occurred in spatially distributed regions across distinct vascular territories, were observed across all tasks, and were also confined to regions of the same functional network (e.g. default mode, dorsal attention) in periods of rest.

### Limitations, remarks and future directions

One potential limitation of our approach is the use of sparsity-promoting L_1_-norm regularized estimators such as LASSO for deconvolution. Even though we observed that sparse 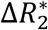-events are sufficient to achieve a high correlation between the preprocessed denoised signals and the BOLD signals estimated with ME-SPFM (see Figure 7), this assumption might not be appropriate for prolonged blocked stimuli or more dense event-related paradigms. In such cases, the proposed ME-based deconvolution framework could be adapted to use other regularization terms, such as L_2_-norm based ridge regression or total variation. Spatial regularization terms could also be added into the algorithm (e.g. following Karahanoğlu et al., 2013) to enhance the spatial robustness of the estimates. We will address the implementation of these approaches in future studies. Importantly, the proposed ME-based deconvolution method can be modified straightforwardly to estimate both 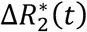 and 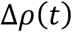 (i.e. also estimating time-varying changes in the net magnetization Δ*S*_0_(*t*)), even using different types of regularization for each component so as to adapt to the nature of their fluctuations (Caballero-Gaudes et al., 2018a; 2018b).

A second limitation of the PFM framework is that it uses a particular HRF shape as the model for deconvolution. Nevertheless, since ME-SPFM is not locked to the timing of the trials, it can clearly account for variability in the onset of the response. It can also describe more complex patterns, such as transient stimulus onset/offset responses (Gonzalez-Castillo et al., 2012) in terms of two 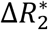-events. Further flexibility can be incorporated by means of structured deconvolution based on multiple basis functions (Caballero-Gaudes et al., 2012).

A third potential limitation is the assumption of linear dependence on TE of fractional BOLD signal changes based on a mono-exponential decay model of the gradient-echo fMRI signal that ME-SPFM builds upon. A biexponential model will be more accurate in voxels with large partial volume effects due to cerebrospinal fluid if the acquisition includes TEs considerably longer than the 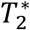 of grey matter (Speck et al., 2001). Furthermore, Havlicek et al. (2017) observed a nonlinear dependence on TE in the human visual cortex at 3T using a combination of multi-echo BOLD and cerebral blood flow measurements, where the amount of nonlinearity varies during the course of the BOLD response, suggesting different physiologically-driven dynamics for the intercept time-course and the BOLD component of the signal. Similarly, Kang et al. (2018) also found a nonlinear dependence on TE mainly in the intravascular signal due to changes in the average chemical exchange time during the stimulation, whereas fractional changes in the extravascular signal can be assumed to linearly increase with TE (Donahue et al., 2011). Yet, these non-linear findings were obtained with long stimulus durations (e.g. visual stimuli of 55 s duration were used in Havlicek et al. (2017) and 16 s duration in Donahue et al. (2011) and Kang et al. (2018)), whereas our data comprised short stimuli of 4 s duration. Even if the assumption of linearity on TE failed, which would make the model assumed in ME-SPFM and other ME-based approaches such as MEICA slightly imperfect, our results showed large agreement with the results of GLM analyses, proving the viability of a ME-based deconvolution approach.

In this work, we used datasets with a known experimental paradigm for validation of ME-SPFM, confirming subject’s compliance with concurrent eye-tracking data. The usage of ME-SPFM in a completely blind scenario with no knowledge of the experimental conditions would be more challenging. This may require the combination of the deconvolution with reverse inference approaches that attempt to decode the subject’s engagement in a particular cognitive process from the activation maps (Poldrack, 2011; Poldrack and Yarkoni, 2016), for example by comparing the activation maps to a predefined set of meta-maps formed using the Activation Likelihood Estimation method of the BrainMap database (Tan et al., 2017). Decoding could be performed at the same rate as the TR of the acquisition, even though successive spatial maps can be averaged to reduce the level of noise in the activation maps and uncertainty in the decoding scores.

Finally, deconvolution algorithms can also be understood as a way of denoising the fMRI signal, like a filtering process matched to the shape of the HRF, wherein the denoised signal comprises the BOLD fluctuations triggered by the deconvolved activity-inducing signal (here, 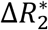 estimates). By some means, this interpretation is supported by the correlation maps of Figure 7 that illustrate a very high correlation between the ME-SPFM fitted dataset, without MEICA, and the preprocessed DN dataset. A comprehensive comparison of ME-SPFM with other ME-based denoising approaches, such as dual-echo regression (Bright and Murphy, 2013) or MEICA-based approaches (Kundu et al., 2012; Power et al., 2018), for denoising the fMRI signal in resting-state and task-based paradigms is beyond the scope of this study. The application of ME-SPFM for denoising will likely involve refinement of the proposed methods (e.g. degree of sparsity, choice of regularization parameters) particularly for connectivity-based analyses.

In summary, in this paper we have introduced the algorithm of multi-echo sparse paradigm free mapping (ME-SPFM) for the deconvolution of BOLD fMRI data collected with ME acquisitions. The ME-SPFM method obtains estimates of the 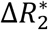 associated with single-trial BOLD events, outperforming our previous method for single-echo acquisitions (1E-SPFM), and exhibiting more concordance with the maps obtained with conventional GLM-based analyses despite being unaware of the timings of the events (i.e. blind detection). The new algorithm is available in AFNI as 3dMEPFM.

## ACKNOWLEDGEMENTS

We thank Prof. Penny A. Gowland for helpful discussion regarding the contents of this manuscript. This research was possible thanks to the support of the Spanish Ministry of Economy and Competitiveness through the Juan de la Cierva Fellowship (IJCI-2014-20821) and Ramon y Cajal Fellowship (RYC-2017-21845), the “Severo Ochoa” Programme for Centres/Units of Excellence in R&D (SEV-2015-490), the National Institute of Mental Health Intramural Research Program (NIH clinical protocol number NCT00001360, protocol ID 93-M-0170, Annual report ZIAMH002783-16), the European Union’s Horizon 2020 research and innovation programme under the Marie Skłodowska-Curie grant agreement No. 713673, and a fellowship from La Caixa Foundation (ID 100010434) (fellowship code LCF/BQ/IN17/11620063). Portions of this study used the high-performance computational capabilities of the NIH High Performance Cluster (Biowulf) at the National Institutes Health, Bethesda, MD (http://hpc.nih.gov).

## SUPPLEMENTARY MATERIALS

**Supplementary material.** Table with the percentage of voxels with 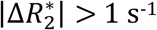 estimated with ME-SPFM and MEICA+ME-SPFM for all datasets, experimental conditions and corresponding trials.

